# Modifiers and Mediators of Craniosynostosis Severity Revealed by Differential Gene Expression

**DOI:** 10.1101/2020.01.28.923508

**Authors:** Amel Dudakovic, Hwa Kyung Nam, Andre vanWijnen, Nan E. Hatch

## Abstract

Severity of craniosynostosis in humans varies widely even in patients with identical genetic mutations. In this study we compared RNA sequencing data from cranial tissues of a severe form of Crouzon craniosynostosis syndrome (C57BL/6 FGFR2^C342Y/+^ mice) with those of a less severe form of Crouzon craniosynostosis (BALB/c FGFR2^C342Y/+^ mice) to identify genetic modifiers that influence craniosynostosis phenotype severity. Comparison of the mice revealed neonatal onset of coronal suture fusion in the form of suture obliteration in C57BL/6 mice (88% incidence, p<.001 between genotypes). Coronal suture fusion in the form of point fusions across the suture occurred at approximately 4 weeks after birth, with less severe skull shape abnormalities, in BALB/c mice. Substantially fewer genes were differentially expressed in BALB/c FGFR2^+/+^ vs. FGFR2C^342Y/+^ mice (87 out of 15,893 expressed genes) than C57BL/6 FGFR2C^+/+^ vs. FGFR2C^342Y/+^ mice (2,043 out of 19,097 expressed genes). Further investigation revealed differential expression of coronal suture fusion associated genes, eph/ephrin boundary genes, cell proliferation genes, osteoblast differentiation genes and epigenetic regulators, among others. The most striking pattern in the data was the minimal change in gene expression seen for most genes in BALB/c FGFR2^+/+^ vs. FGFR2C^342Y/+^ mice. Analysis of protein processing and lysosomal components support the hypothesis that the craniosynostosis phenotype is less severe in BALB/c mice because the mutant FGFR2C^342Y^ protein is not expressed to the same extent as that seen in C57BL/6 mice. Together, these results suggest that a strategy aimed at increasing degradation of the mutant receptor or downstream signaling inhibition could lead to diminished phenotype severity.

## Introduction

Craniosynostosis is the pediatric condition of premature cranial bone fusion that occurs in approximately 1 per 2000 live births (McCarthy et al., 2012). This condition can lead to high intracranial pressure, abnormal skull and facial shapes, blindness, seizures and brain abnormalities (Flapper et al., 2009; McCarthy et al., 2012; Morriss-Kay and Wilkie, 2005; Okajima et al., 1999; Rasmussen et al., 2008; Seruya et al., 2011). Because the sole treatment is cranial surgery, even with appropriately early diagnosis severely affected patients can suffer high medical and financial morbidity (Abe et al., 1985; Baird et al., 2012; Renier et al., 2000). Surgical approaches also do not fully correct abnormal skull and facial shapes, which contribute to psychosocial challenges.

Craniosynostosis can occur in isolation or in association with over 105 different genetic syndromes and 174 identified genetic mutations (OMIM, 2019). The pathogenesis of craniosynostosis can include abnormal mesenchymal progenitor cell migration to sites of bone and suture formation, cranial bone and suture boundary formation/maintenance, premature cranial progenitor cell lineage commitment and/or diminished proliferation of cranial bone and suture cells (Chen et al., 2003; Deckelbaum et al., 2012; Eswarakumar et al., 2004; Hayano et al., 2015; Holmes et al., 2009; Komatsu et al., 2013; Liu et al., 2013; Liu et al., 2015; Merrill et al., 2006; Rice et al., 2010; Teng et al., 2018; Yousfi et al., 2002). Mechanisms differ depending upon the genetic basis (known or unknown) and involved suture(s).

Crouzon craniosynostosis occurs due to inherited or germline somatic activating mutations in the mesenchymal splice form of *Fgfr2 (FGFR2IIIc)* (Chen et al., 2003; Eswarakumar et al., 2004; Reardon et al., 1994; Wilkie et al., 1995). Crouzon syndrome has been estimated to represent 4.8% of craniosynostosis cases at birth (Cohen and Kreiborg, 1992). It is a commonly held belief that the craniosynostosis associated FGFR mutations act as gain of function mutations in terms of FGF signaling. More specifically, Crouzon syndrome linked mutations in FGFR2IIIc can result in ligand independent autophosphorylation, dimerization and tyrosine kinase activity in vitro (Galvin et al., 1996; Neilson and Friesel, 1995; Robertson et al., 1998). Notably, individuals with Crouzon craniosynostosis exhibit variable phenotype expression, even when carrying identical mutations (Lajeunie et al., 2000; Sher et al., 2008). Those with a more severe phenotype experience greater morbidity. The objective of this study was to identify potential modifiers and mediators of craniosynostosis phenotype severity by gene expression analysis of tissues from the FGFR2^C342Y/+^ mouse model of Crouzon syndrome on two different genetic background strains, one of which exhibits a severe craniosynostosis phenotype and another which exhibits a mild craniosynostosis phenotype.

## Materials and methods

### Animals and Phenotyping

FGFR2^C342Y/+^mice were backcrossed with C57BL/6 and BALB/c mice (obtained from Charles River Laboratories) for at least fifteen generations prior to experiments. C57BL/6 mice have a severe phenotype with coronal suture fusion typically initiating in neonatal mice. BALB/c FGFR2^C342Y/+^mice have a more moderate form phenotype with craniosynostosis in the form of small point fusions across the coronal suture first apparent four weeks after birth (Liu et al., 2013). Genotyping was performed as previously described (Eswarakumar et al., 2004; Liu et al., 2013). Briefly, DNA from tail digests was amplified by polymerase chain reaction using 5’-gagtaccatgctgactgcatgc-3’and 5’-ggagaggcatctctgtttcaagacc-3’ primers to yield a 200 base pair band for wild type FGFR2 and a 300 base-pair band for mutant FGFR2^C342Y^. Mice were fed standard powdered rat chow (Harlan Laboratories) and distilled water *ad libitum*, and housed under standard 12 hour dark/light cycles. Genders were combined for analyses. Neonatal C57BL/6 FGFR2^+/+^ (n=3), C57BL/6 FGFR2^C342Y/+^ mice (n=3), BALB/c FGFR2^+/+^ (n=3) and BALB/c FGFR2^C342Y/+^ (n=3) skulls were dissected to collect the frontal bone with nasal bone, frontonasal suture and coronal sutures for RNA sequencing from each of the mice. For phenotyping, three-week-old BALB/c mice (n=13 FGFR2^+/+^, n=11 FGFR2^C342Y/+^) and C57BL/6 mice (n=14 FGFR2^+/+^, n=17 FGFR2^C342Y/+^) were euthanized by CO2 overdose for micro CT and for measurements of height and weight. Animal use followed federal guidelines and were performed in accordance with the University of Michigan’s Institutional Animal Care and Use Committee. The primary phenotype outcome assessments were coronal suture fusion incidence and differences in craniofacial skeletal shape. Secondary outcome assessments were incidence of malocclusion and cranial bone density measurements.

### Micro CT

Whole dissected skulls from three-week-old mice were fixed then scanned in water at an 18 μm isotropic voxel resolution using the eXplore Locus SP micro-computed tomography imaging system (GE Healthcare Pre-Clinical Imaging, London, ON, Canada). Measurements were taken at an operating voltage of 80 kV and 80 mA of current, with an exposure time of 1600 ms using the Parker method scan technique (Parker, 1982), which rotates the sample 180 degrees plus a fan angle of 20 degrees.

### Cranial Suture Assessment

Fusion of the coronal suture was identified on micro CT scans of dissected mouse skulls. The coronal suture was viewed using the two-dimensional micro CT slices in an orthogonal view across the entire length of the suture, as previously described (Liu et al., 2013; Liu et al., 2014).

### Linear Measurements

Craniofacial linear skeletal measurements were taken using digital calipers on dissected skulls using previously reported craniofacial skeletal landmarks (Liu et al., 2013; Perlyn et al., 2006; Richtsmeier et al., 2000), including standard measurements currently in use by the Craniofacial Mutant Mouse Resource of Jackson Laboratory (Bar Harbor, ME). Linear measurements were normalized to total skull length (measured from nasale to opisthion) to account for size differences between background strains as well as between FGFR2^+/+^ and FGFR2^C342Y/+^ mice. Please note that two different skull length measurements were taken such that even skull length is reported as normalized for skull size. Measurements were performed twice and an average of the two measurements was utilized for statistical comparison by genotype and treatment.

### Phenotype Statistics

Descriptive statistics (mean, standard deviation) for each parameter were calculated for all phenotype measurements. Student’s t-test was used for comparison between groups for craniofacial linear measurements. Fisher’s exact test was used to compared the incidence of coronal suture fusion and the incidence of malocclusion between genotypes and background strains.

### RNA Isolation

Dissected postnatal day 3 cranial tissues were rinsed then digested (2mg/ml collagenase type II + 0.05% trypsin in aMEM). After digestion, the frontal bone (including nasal bone, coronal and frontonasal sutures) tissues were separated from parietal bone tissues, placed into microcentrifuge tubes with Zirconium Oxide beads (Next Advance) and Trizol (Invitrogen), then homogenized according to the manufacturer protocol of Bullet Blender (BBY24M, Next Advance). After homogenization, RNA was extracted from the supernatant using Direct-zol RNA MiniPrep (Zymo Research). Extracted RNA was diluted in DNase/RNase-free water and stored at −80C until use.

### RNA sequencing

RNA sequencing was performed as previously reported (Paradise et al., 2019). Expression values for each gene were normalized to one million reads and corrected for gene length (reads per kilobase per million mapped reads, RPKM). RPKM values were determined for each genotype and strain group using thee independent biologic replicates.

Quality of the raw reads data for each sample was checked using FastQC version v0.11.3. We used default parameter settings for alignment, with the exception of: “b2-very-sensitive” telling the software to spend extra time searching for valid alignments. We used FastQC for a second round of quality control (post-alignment), to ensure that only high quality data would be input to expression quantitation and differential expression analysis. We used HTSeq version 0.6.1/DESeq2 version 1.14.1 (Zhang et al., 2014), using UCSC mm10.fa as the reference genome sequence (Kent WJ, 2002).

We identified genes and transcripts as being differentially expressed based on three criteria: test status = “OK”, FDR ≤ 0.05, and fold change ≥ ± 1.5. Expression quantitation was performed with HTSeq version 1.14.1, to count non-ambiguously mapped reads only. Data were prefiltered to remove genes with 0 counts in all samples. Normalization and differential expression were performed with DESeq2 using a negative binomial generalized linear model. Principal component analysis and non-normalized distribution plots indicated good quality data consistent with the set of differentially expressed genes. Samples did not appear to strongly cluster by background genotype which were divided between batches. Therefore, batch effect is unlikely to have a large impact.

The data was analyzed using Advaita Bio’s iPathway software (https://www.advaitabio.com/ipathwayguide), which implements the ‘Impact Analysis’ approach, as described in (Draghici, 2007, Tarca, 2008, Donato, 2013, Ahsan 2018).” The dataset is available at the Gene Expression Omnibus, accession number GSE143628. Comparisons between groups (three independent biologic replicates per group) were made using post hoc student’s t-tests and a Bonferroni correction for multiple comparisons. Statistical significance was established as p<0.05 (p<.0125 with Bonferroni correction). Data is presented in metaanalysis format, with each graph showing differential gene expression for four comparisons: 1) C57BL/6 FGFR2^+/+^ vs. BALB/c FGFR2^+/+^ cranial tissues, 2) C57BL/6 FGFR2^C342Y/+^ vs. BALB/c FGFR2^C342/+^ cranial tissues, 3) C57BL/6 FGFR2^C342Y/+^ vs. C57BL/6 FGFR2^+/+^ cranial tissues, and 4) BALB/c FGFR2^C342Y/+^ vs. BALB/c FGFR2^+/+^ cranial tissues.

### Real Time PCR

RNA isolated for RNA sequencing was used to perform real time polymerase chain reaction. mRNA levels were assayed by reverse transcription and real time PCR. Real time PCR was performed for murine Gapd, Padi2, Gata1, E2F2 and Uchl1 using Taman primer sets and Taqman Universal PCR Master Mix (Applied Biosystems). Real-time PCR was performed on a 7500 Real Time PCR system (Applied Biosystems) and quantified by comparison to a standard curve.

## Results

### Craniofacial Phenotype is Severe in C57BL/6 but not in BALB/c Crouzon Mice

#### Craniofacial Shape

Representative isosurface micro CT images of skulls demonstrate that Crouzon mice carrying the FGFR2^+/C342Y^ mutation differ in morphology from their wild type counterparts, and that the shape differences between mutant and wild type are more apparent in mice on the C57BL/6 than the BALB/c background (**Fig. 1**). C57BL/6 FGFR2^+/C342Y^ skulls are tall, dome shaped and exhibit midface hypoplasia that is more severe than that seen in BALB/c FGFR2^+/C342Y^ mice. Craniofacial skeletal linear measurements normalized to total skull length confirmed the consistencies of the above observations and revealed many differences between FGFR2^342Y/+^ Crouzon and FGFR2^+/+^ wild type mice in terms of skull shape on both congenic backgrounds (**Fig. 2**). Between the two strains, C57BL/6 FGFR2^342Y/+^ Crouzon mice had greater changes in skull width, nose length and inner canthal distances when compared to BALB/c FGFR2^342Y/+^ Crouzon mice.

**Figure 1.**
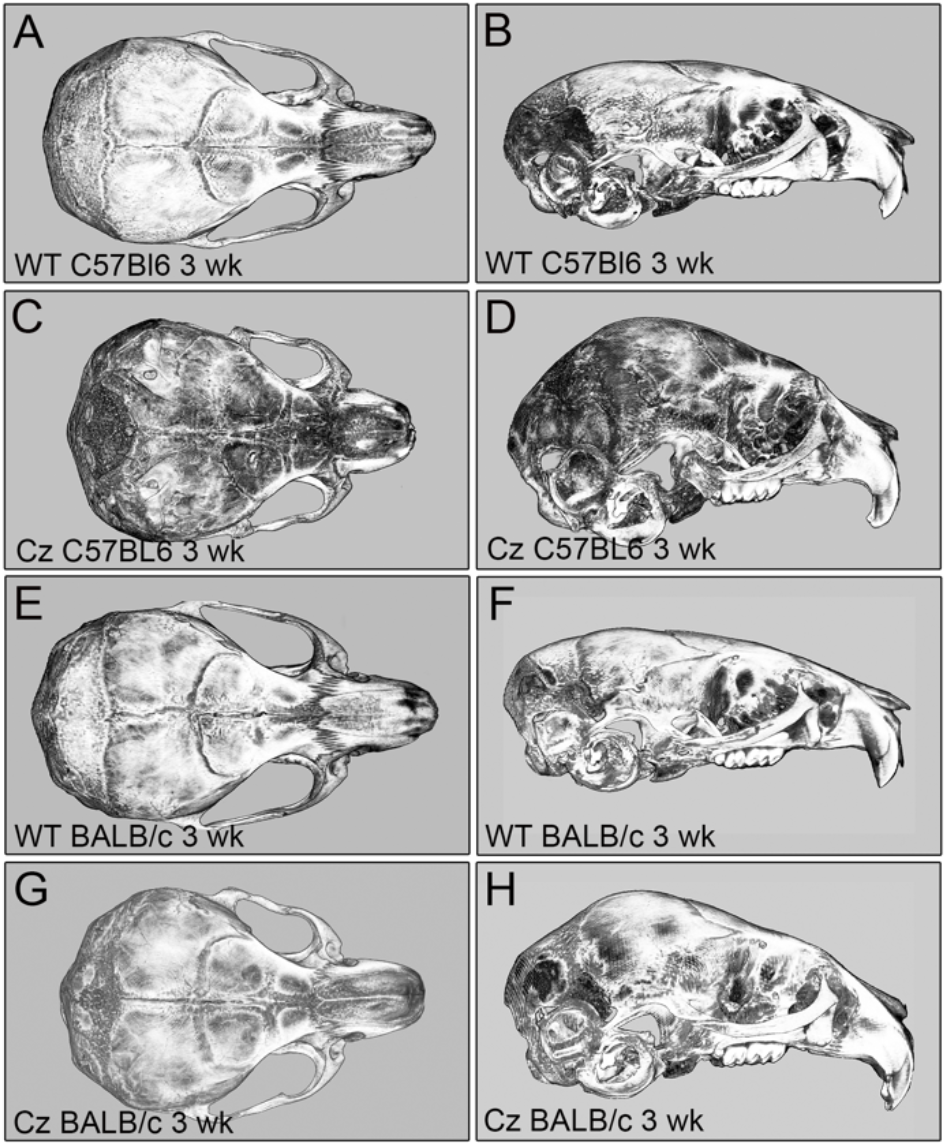
Micro CT images of C57BL/6 and BALB/c FGFR2^+/+^ and FGFR2^C342Y/+^ mice. Micro CT isosurface images of day 21 FGFR2^C342Y/+^ Crouzon (CZ) and wild type (WT) mice on the C57BL/6 and BALB/c congenic backgrounds are shown in axial view from above (A,C,E,G) and lateral view (B,D,F,H). Note skull shape differences in FGFR2^+/+^ vs. FGFR2^C342Y/+^ mice that are more severe on the C57BL/6 background. Darker bone is bone of diminished density.

### Craniosynostosis and Malocclusion

Analysis of craniosynostosis revealed a high incidence of coronal suture fusion in three-week-old C57BL/6 FGFR2^C342Y/+^ mice as compared to FGFR2^+/+^ mice (88% vs. 0%, p<.001). In contrast, the coronal suture was not fused in any in three-week-old BALB/c FGFR2^+/+^ or FGFR2^C342Y/+^ mice. A class III malocclusion was evident in 86% of C57BL/6 FGFR2^C342Y/+^ mice and in 27% of BALB/c FGFR2^C342Y/+^ (p<.03), while no malocclusion was identified in FGFR2^+/+^ mice on either genetic background.

### Identification of Differentially Expressed Genes in C57BL/6 and BALB/c Cranial Bones

Overall gene expression analyses showed that for some individual comparisons, there are over 1,000 differentially expressed genes, while other comparisons have less than 10 unique differentially expressed genes that reached statistical significance (**Fig. 3**). To look for preexisting modifiers of differentially expressed genes between the two genetic strains, we compared RNA from FGFR2^+/+^ C57BL/6 and BALB/c cranial tissues. To look for primary mediators of craniosynostosis upon expression of the mutated receptor that are genetic background dependent, we compared RNA from FGFR2^+/+^ C57BL/6 with FGFR2^+/+^ BALB/c tissues, FGFR2^C342Y/+^ C57BL/6 with FGFR2^C342Y/+^ BALB/c tissues, and compared FGFR2^+/+^ with FGFR2^C342Y/+^ tissues in C57BL/6 and BALB/c strains. Volcano plots demonstrate differential gene expression between C57BL/6 and BALB/c FGFR2^+/+^ tissues, and between C57BL/6 and BALB/c FGFR2^C342Y/+^ tissues (**Fig. 4**). Of note, substantially fewer genes were differentially expressed in BALB/c FGFR2C^342Y/+^ vs. FGFR2^+/+^ mice (87 out of 15,893 genes with measured expression) than in C57BL/6 FGFR2C^342Y/+^ vs. FGFR2^+/+^ mice (2,043 out of 19,097 genes with measured expression). When limiting to those genes that were only differentially expressed in the individual comparisons (FGFR2C^342Y/+^ vs. FGFR2^+/+^ tissues on the two different strains), 4 genes were found differentially expressed between genotypes on the BALB/c strain while 409 were found differentially expressed between genotypes on the C57BL6 strain.

**Figure 2.**
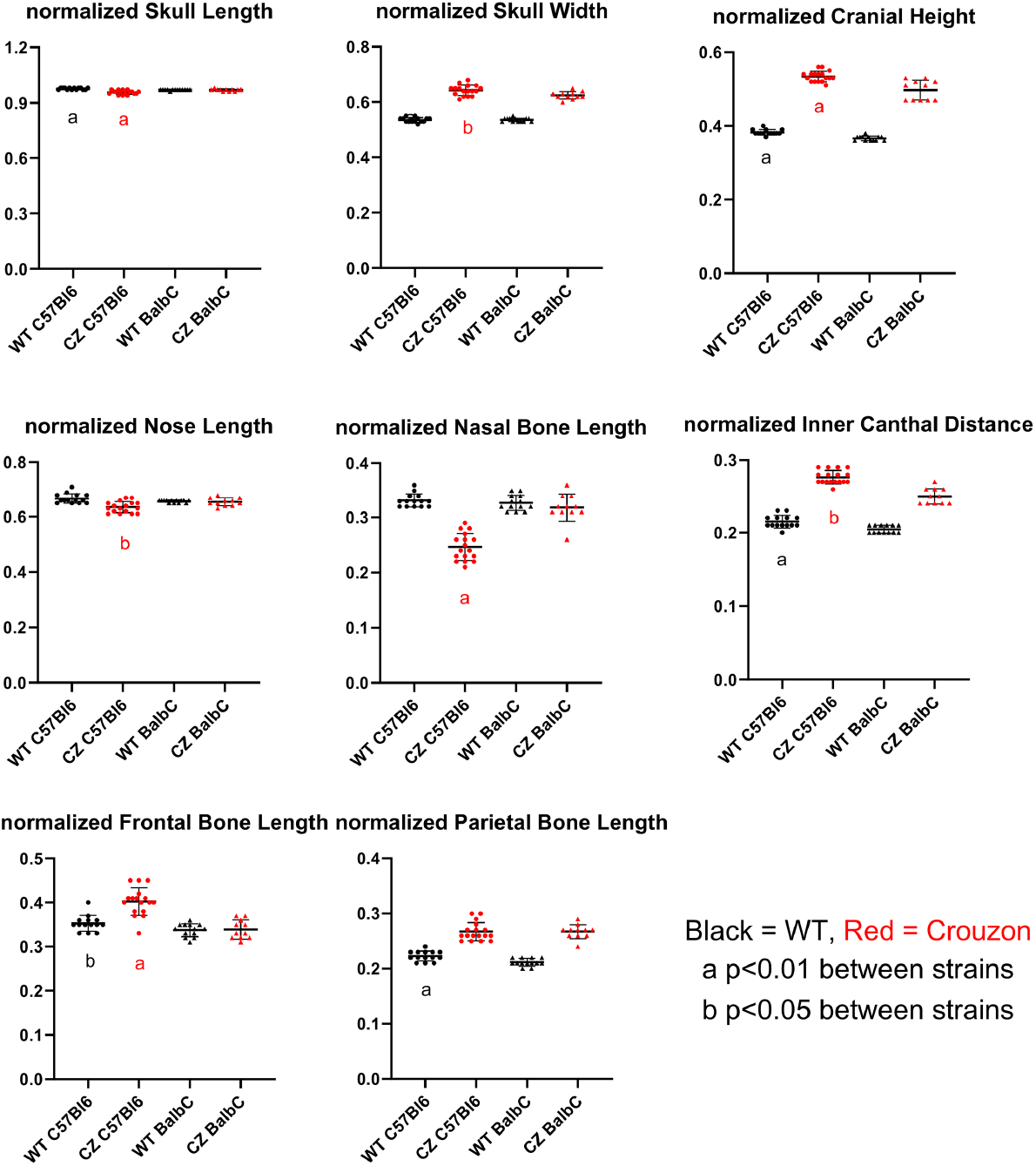
Craniofacial linear measurements of C57BL/6 and BALB/c FGFR2^+/+^ and FGFR2^C342Y/+^ mice. Craniofacial linear measurements are presented as normalized to total skull length (measured from nasale to opisthion). The skull length measurement presented here is from nasale to paro (intersection of interparietal and anterior aspect of occipital bones at the midline). Note that the C57BL/6 FGFR2^342Y/+^ Crouzon mice had greater changes in skull width, nose length and inner canthal distances when compared to BALB/c FGFR2^342Y/+^ Crouzon mice, indicative of an overall more brachycephalic and acrocephalic skull shape. Data from n=13 BALB/c FGFR2^+/+^, n=11 BALB/c FGFR2^C342Y/+^, n=14 C57BL/6 FGFR2^+/+^ and n=17 C57BL/6 FGFR2^C342Y/+^ mice are shown.

**Figure 3.**
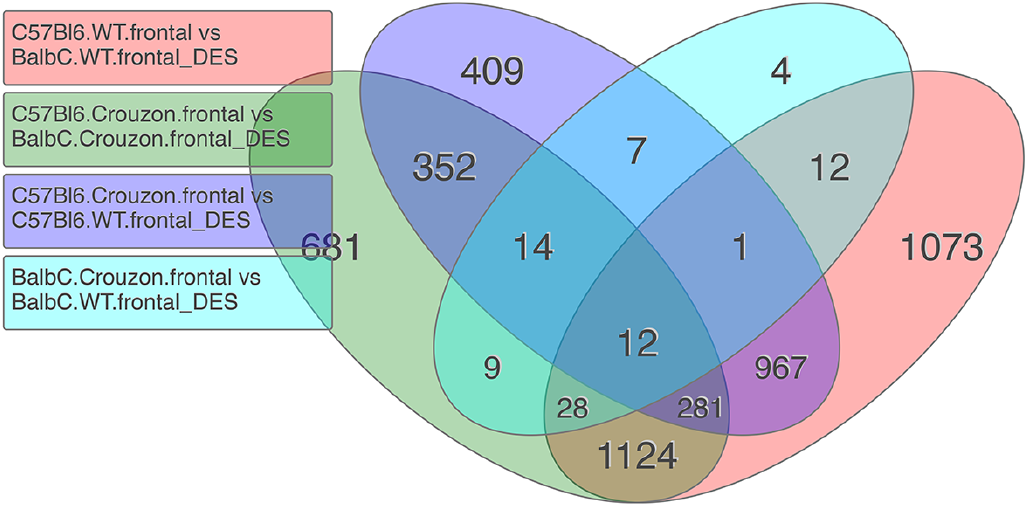
Differential Gene Expression in neonatal FGFR2^+/+^ and FGFR2^C342Y/+^ mice on C57Bl/6 and Balb/C inbred genetic strains. Venn diagram presentation of differentially expressed genes from three independent biologic replicates per group demonstrates 1073 uniquely differentially expressed genes in C57BL/6 vs. BALB/c FGFR2^+/+^ cranial tissues, 681 uniquely differentially expressed genes in C57BlL6 vs. BALB/c FGFR2^C342Y/+^ cranial tissues, 409 uniquely differentially expressed genes in C57BL/6 FGFR2^+/+^ vs. FGFR2^C342Y/+^ cranial tissues and 4 uniquely differentially expressed genes in BALB/c FGFR2^+/+^ vs. FGFR2^C342Y/+^ cranial tissues. N=3 independent biologic replicates per group.

**Figure 4.**
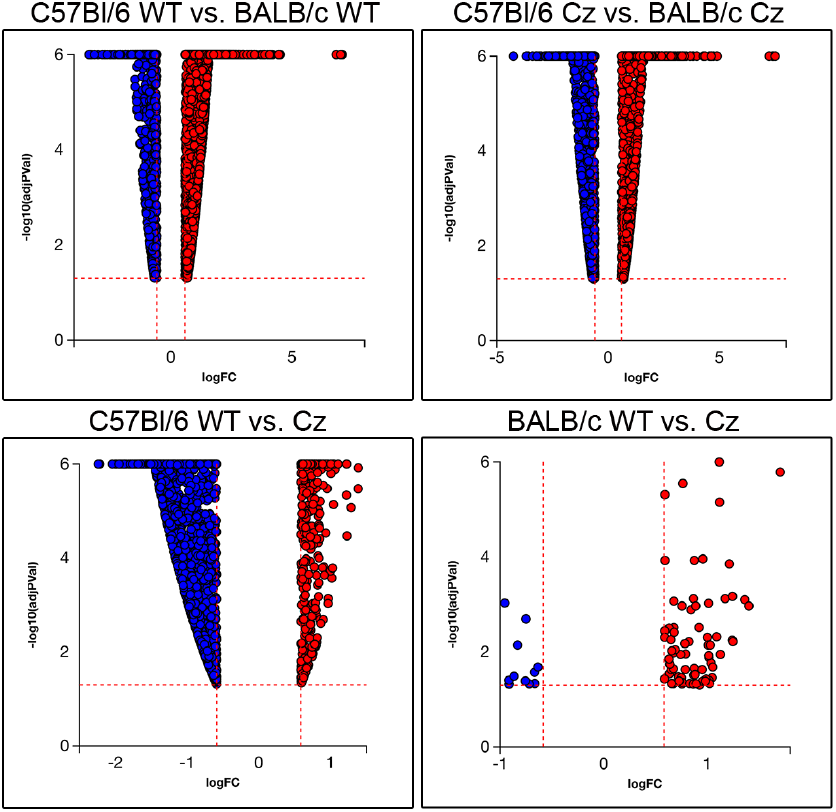
Volcano plots demonstrate differential gene expression by magnitude and statistical significance for the individual comparisons. Note that a greater number of genes are differentially expressed and at a higher magnitude of differential expression in C57BL/6 FGFR2C^342Y/+^ vs. FGFR2^+/+^ tissues than BALB/c FGFR2C^342Y/+^ vs. FGFR2^+/+^ tissues. N=3 independent biologic replicates per group.

### Differentially Expressed Craniosynostosis Associated Genes

We first searched for genes under the GO term craniosynostosis (*Msx1, Msx2, Twistl, Tcfl2, Alx4, Erf, Zic1*). Because Crouzon syndrome in humans is most commonly associated with fusion of coronal as opposed to the sagittal suture, and because the FGFR2^C342Y/+^ mouse model of Crouzon syndrome on C57BL6 and BALB/c backgrounds exhibits coronal but not sagittal suture fusion, we also looked for differential expression of genes associated with coronal suture fusion including *Fgfr1, Fgfr2, Fgfr3, Efnb1, EphA4, En1 and Ezh2* (**Fig. 5**). Differential RNA expression was not found for *Fgfr1, Fgfr2, Fgfr3, Efnb1, EphA4, Tcf12, Erf or Alx4*. Results showed significantly downregulated expression of *Msx1, Msx2, Twist1* and *Zic1* in C57BL/6 as compared to BALB/c FGFR2^+/+^ mice, and downregulated expression of *Msx1, Twist2* and *Zic1* in C57BL/6 as compared to BALB/c FGFR2^C342Y/+^ mice. No difference in *Msx2* or *Twist1* gene expression was found between C57BL/6 FGFR2^C342Y/+^ and BALB/c FGFR2^C342Y/+^ mice. Notably, expression of *Msx2, Zic1 and Twist1* expression was increased in C57BL/6 FGFR2^C342Y/+^ as compared to C57BL/6 FGFR2^+/+^ mice while no differences were noted between BALB/c FGFR2^C342Y/+^ as compared to BALB/c FGFR2^+/+^ mice.

**Figure 5.**
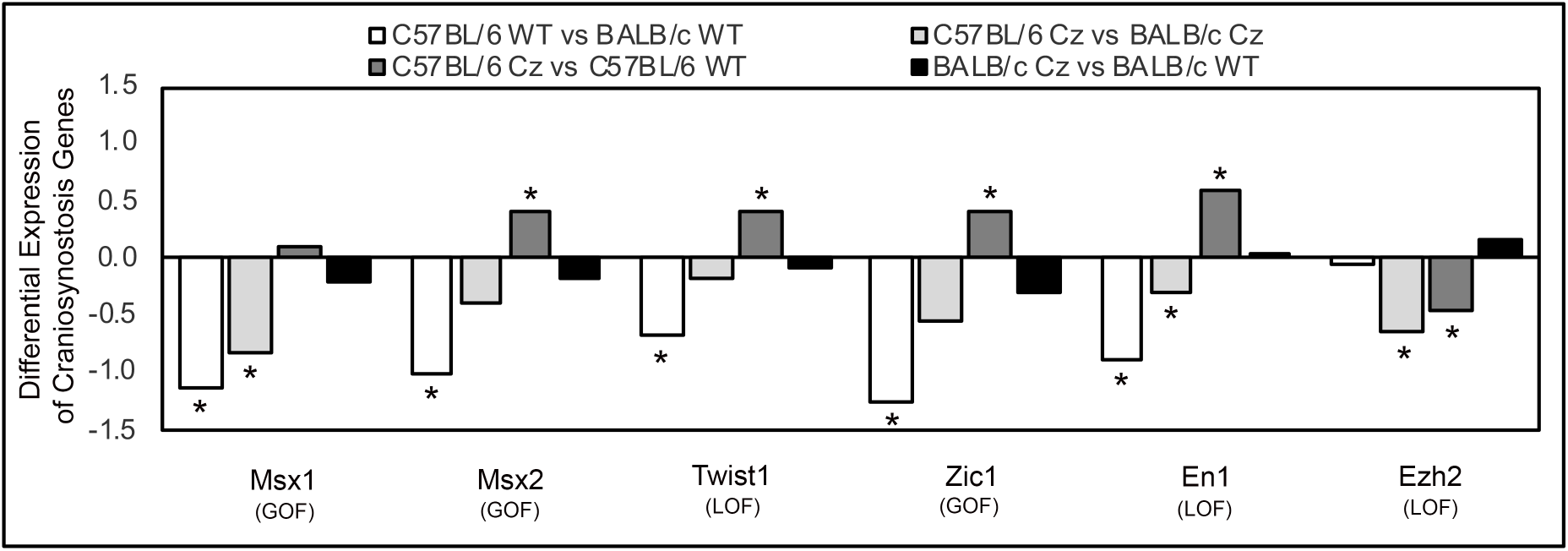
Differential Expression of Craniosynostosis Associated Genes. Differentially expressed genes for the GO term craniosynostosis plus other genes known to be associated with craniosynostosis in the form of coronal suture fusion are shown. Data for each gene is presented for four different individual comparisons, as indicated by color. Differential expression was not found for *Fgfr1, Fgfr2, Fgfr3, Efnb1, EphA4, Tcf12, Erf or Alx4*. N=3 independent biologic replicates per group.

### The Role of Coronal Suture Boundary Formation and Maintenance

We found that *En1* (engrailed 1) expression was significantly diminished in C57BL/6 compared to BALB/c cranial tissues regardless of mutation (FGFR2^+/+^ or FGFR2^342Y/+^) (**Fig. 5**). Notably, *En1* expression was not different between mutant and wild type mice on the BALB/c background. In contrast, *En1* was expressed to a significantly greater extent in C57BL/6 FGFR2^342Y/+^ as compared to C57BL/6 FGFR2^+/+^ cranial tissues indicating that expression of the mutant receptor feeds back on *En1* expression in C57BL/6 mice. *En1* is a homeodomain transcription factor previously shown to be essential for proper mesoderm/neural crest boundary formation of the coronal suture (Deckelbaum et al., 2012), such that low En1 gene expression levels could predispose to a more severe craniosynostosis phenotype. Consistent with this hypothesis, we also found significant differences in eph and ephrin gene expression by genetic background strain and in mutant vs. wild type C57BL/6 but not BALB/c mice (**Supplemental Fig. 1**).

### Differential Expression of Epigenetic Regulators

Epigenetic changes in chromatin are regulated by enzymes that edit or erase histone post-translational modifications such as phosphorylation, acetylation, methylation and citrullination. We found that the H3K27 methyl transferase *Ezh2* was significantly downregulated upon expression of the FGFR2^C342Y^ mutation in C57BL/6 compared to BALB/c mice (**Fig. 5**). Genetic depletion of *Ezh2* predisposes to coronal suture fusion (Dudakovic et al., 2014), therefore this gene may act as a primary mediator to increase craniosynostosis phenotype severity by altering the chromatin landscape in cranial progenitor cells downstream of the mutant FGFR2^C342Y^ receptor in C57BL/6 mice.

PCA analysis of epigenetic regulators also reveals segregation by genotype and strain (**Supplemental Figure 2**). Analysis of the RNA seq data by the GO term histone modifier genes revealed thirteen histone modifying genes to be differentially expressed between the groups (**Supplemental Table 1**). In addition, a search for GO term nucleosome reveals numerous differences between C57BL/6 and BALB/c cranial tissues (**Supplemental Table 2**) regardless of mutation (FGFR2^+/+^ or FGFR2^342Y/+^). In addition, no nucleosome genes are differentially expressed in BALB/c FGFR2^+/+^ compared to FGFR2^342Y/+^ tissues, while six genes are significantly downregulated by expression of the FGFR2^342Y^ mutant receptor in C57BL/6 mice. Together this data indicates both that the chromosome landscape is different in FGFR2^+/+^ C57BL/6 vs. FGFR2^+/+^ BALB/c mice, and in FGFR2^342Y/+^ C57BL/6 vs. FGFR2^342Y/+^ BALB/c mice. It should also be noted that Padi2, a histone H3-R26 citrullination enzyme known for controlling proliferation and cell cycle progression in a tissue type specific manner (Guo et al., 2017), is within the top 45 upregulated genes when comparing C57BL/6 with BALB/c FGFR2^+/+^ mice (3-fold up in C57BL/6).

### Differential Expression of Protein Modification and Degradation Genes

The pattern of minimal difference in gene expression between mutant and wild type mice on the BALB/c background is noticeable and repeatable in the RNA seq data. This pattern in the data suggested diminished expression and/or activity of the mutant FGFR2^C342Y^ receptor in BALB/c mice, which led us next to look for changes in gene expression that could account for differential receptor processing and/or degradation between the two strains. To investigate gene changes that may lead to diminished mutant receptor protein expression in BALB/c as compared to C57BL/6 mice, we analyzed the sequencing data for differences in genes that could influence wild type and mutant FGFR2 modification and degradation (**Fig. 6**). The N-glycosylation enzymes (*Mgat* and *Mgat5b)* exhibit significantly greater expression in C57BL/6 FGFR2^+/+^ and FGFR2^342Y/+^ as compared to BALB/c FGFR2^+/+^ and FGFR2^342Y/+^ cranial tissues, indicating that the mutant FGFR2^342Y^ protein may have a greater chance of complete glycosylation in the C57BL/6 than the BALB/c mice. ER/Golgi chaperone proteins *Hspa1a* and *Hspa1b* are also both expressed more in C57BL/6 FGFR2^+/+^ and FGFR2^342Y/+^ as compared to BALB/c FGFR2^+/+^ and FGFR2^342Y/+^ cranial tissues. Additionally, analysis of the RNA seq data by the GO term lysosomes revealed significantly decreased expression of peroxidase and protease enzymes *Mpo, Prtn3* and *Prss57* in C57BL/6 FGFR2^342Y/+^ as compared to BALB/c FGFR2^342Y/+^ tissues. It should also be noted that the *Uchl1* gene for ubiquitin carboxy-terminal hydrolase L1 deubiquitinating enzyme is the 41st top upregulated gene in C57BL/6 compared to BALB/c FGFR2^+/+^ tissues (3-fold up in C57BL/6), 51^st^ top upregulated gene in C57BL/6 compared to BALB/c FGFR2^C342Y/+^ tissues (2.6-fold up in C57BL/6).

**Figure 6.**
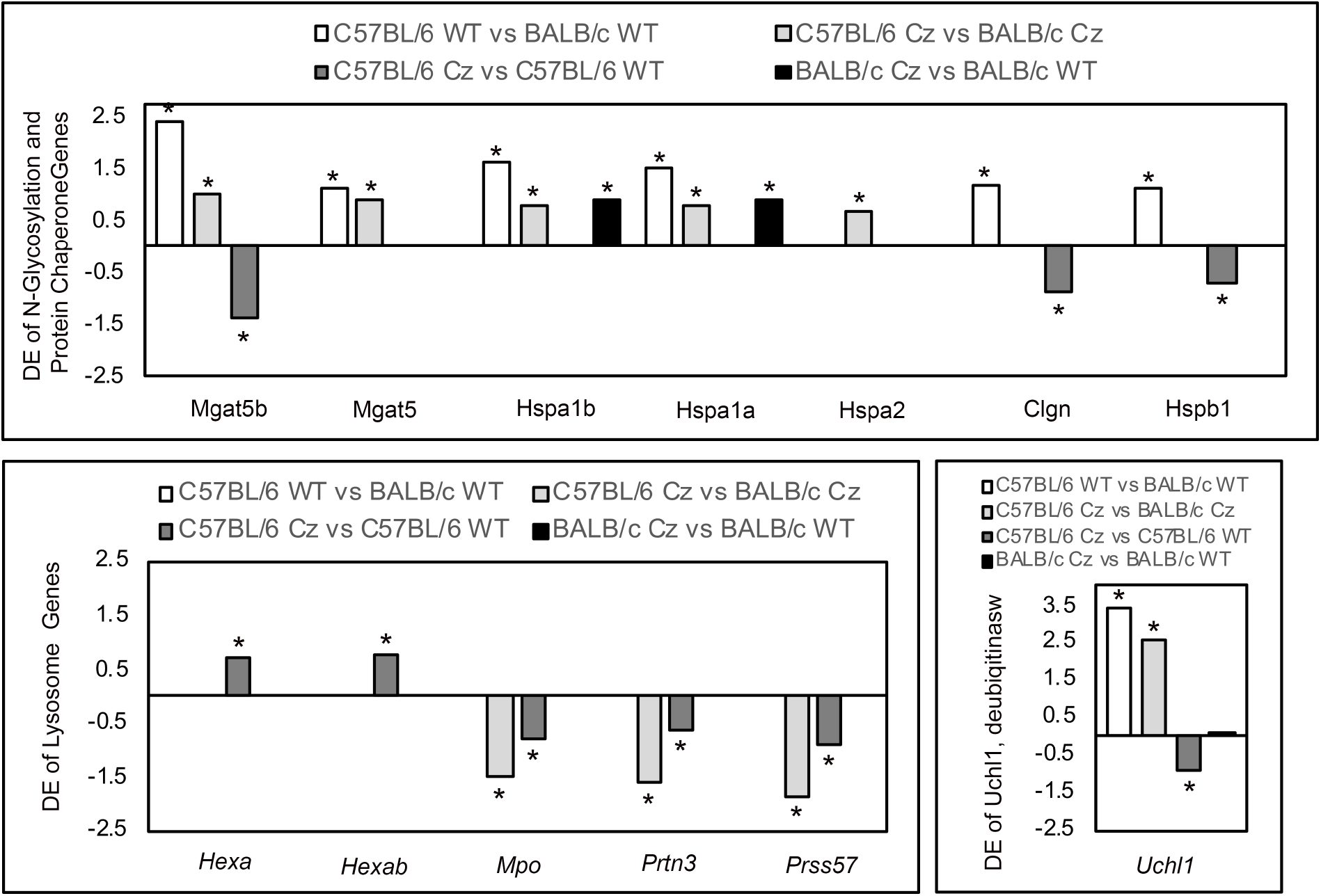
Differential Expression of Protein Processing and Protein Degradation Genes. Differentially expressed genes for N-glycosylation enzymes, ER/Golgi protein chaperones, lysosomal components and the ubiquitin carboxy-terminal hydrolase L1 deubiquitinating enzyme *Uchl1* are shown for each of the individual comparisons. N=3 independent biologic replicates per group.

**Figure 7.**
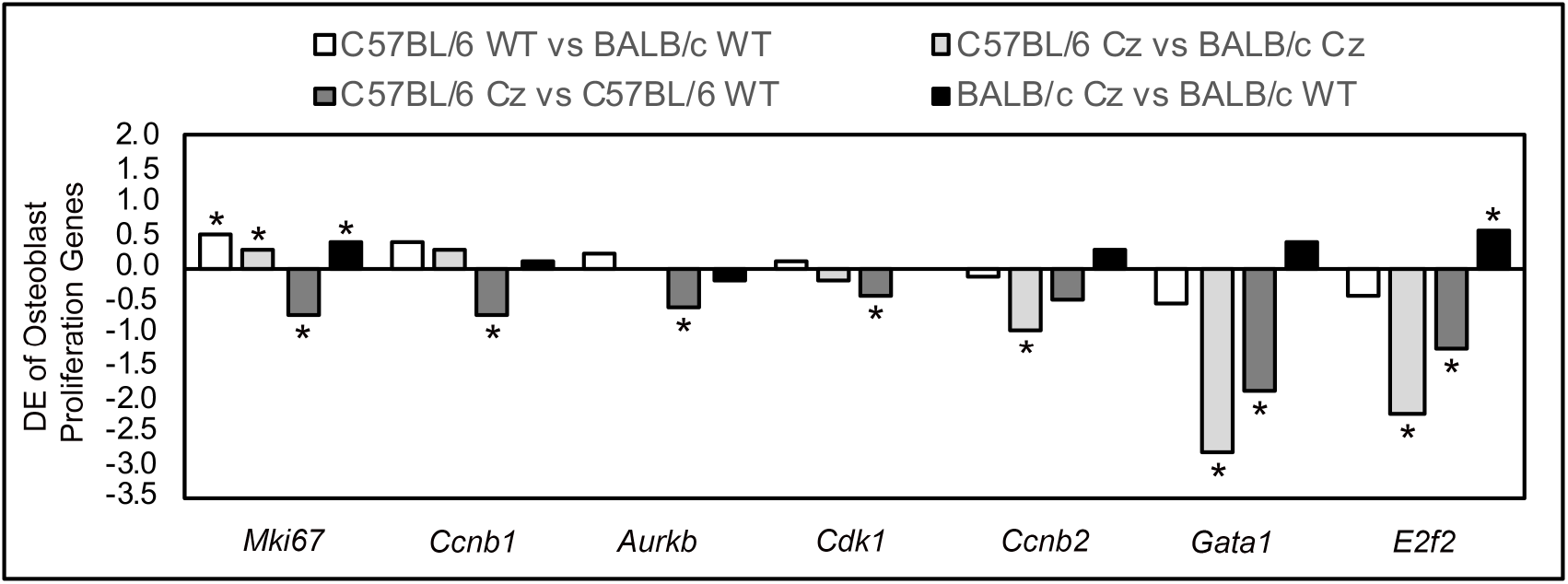
Differential Expression of Osteoblast Proliferation Genes. Differentially expressed genes related to proliferation and cell cycle progression, as well as Gata1 from the GO term positive regulator of osteoblast proliferation are is shown for each of the individual comparisons. N=3 independent biologic replicates per group.

**Figure 8.**
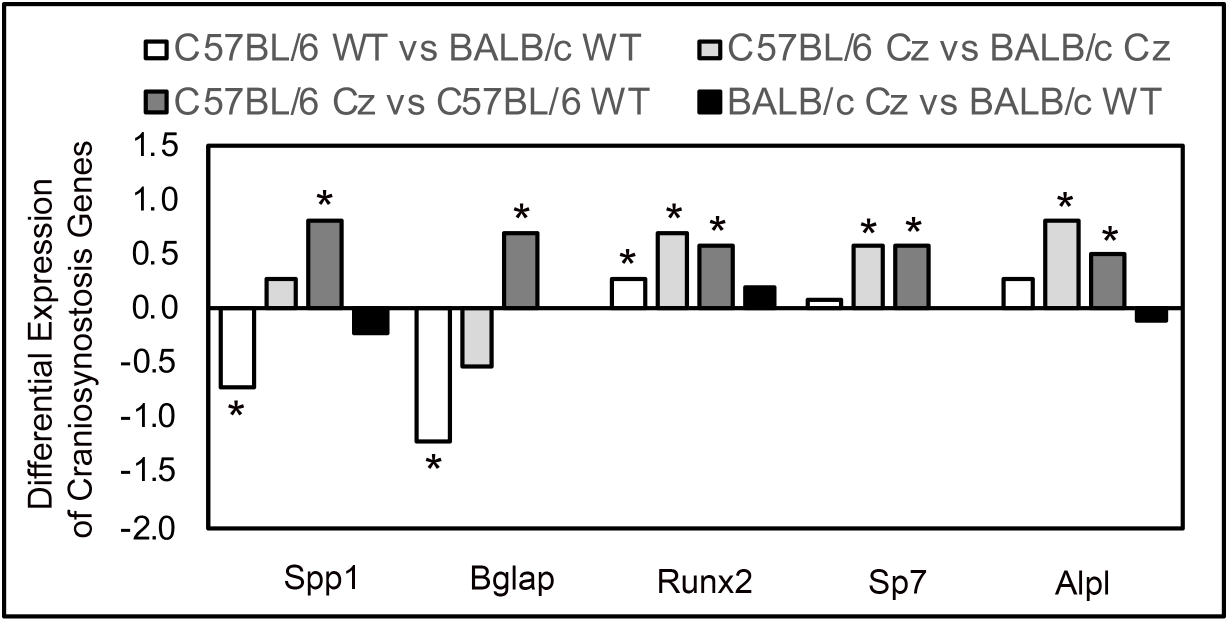
Differential Expression of Osteoblast Differentiation Genes. Differentially expressed genes for markers of osteoblast differentiation are is shown for each of the individual comparisons. N=3 independent biologic replicates per group.

**Figure 9.**
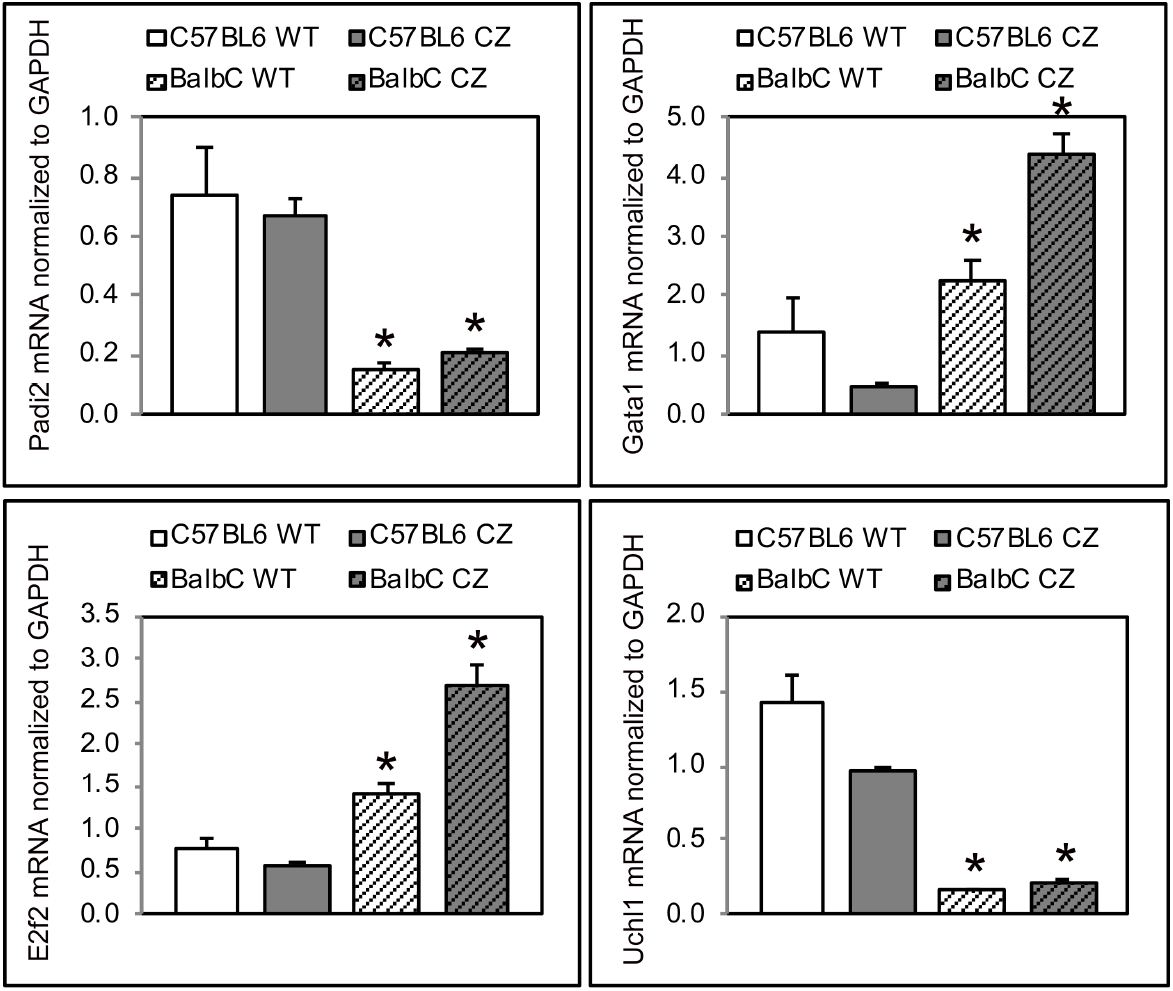
Real Time PCR of Differentially Expressed Genes. RNA was isolated from postnatal day 3 cranial tissues, same as for RNA sequencing analysis. *Padi2, Gata1, E2h2* and *Uchl1* mRNA levels were measured by real time PCR. Results are presented as normalized to GAPDH. *p<.05 between genetic background strains.

### Differential Expression of Osteoblast Proliferation and Differentiation genes

Premature loss of cranial suture tissue due to diminished progenitor cell proliferation and/or premature progenitor osteoblast differentiation are presumed by many investigators to be contributing cellular mechanisms behind craniosynostosis. Therefore, we next analyzed the samples by the GO term proliferation. Results revealed 397/1766 genes differentially expressed between C57BL/6 and BALB/c FGFR2^+/+^ tissues (p<.002), 300/1752 genes between C57BL/6 and BALB/c FGFR2^C342Y/+^ tissues (p<.002), 263/1835 genes between C57BL/6 FGFR2^+/+^ and FGFRC342Y^+/+^ tissues (p<.0002), and 13/1697 genes between BALB/c FGFR2^+/+^ and FGFRC^342Y+/+^ tissues (p not significant). Looking specifically at expression of the proliferation marker *Mki67*, as well as cyclins, cyclin dependent kinases, the mitotic spindle protein Aurkb (Goldenson and Crispino, 2015) and the DNA replication and mitosis promoting E2F2 transcription factor (Ishida et al., 2001; Lees et al., 1993), the most obvious pattern to emerge is that expression of the FGFR^C342Y^ mutant receptor downregulates expression of the majority of proliferation genes that are differentially expressed on in C57BL/6 mice. Notably, E2F2 is the top 55^th^ downregulated gene in C57BL/6 vs. BALB/c FGFR2^C342Y/+^ tissues (2.2-fold down in C57BL/6). Differential expression of the transcription factor Gata1 from the GO term osteoblast positive regulator of proliferation is also shown as downregulated in C57BL/6 vs. BALB/c FGFR2^C342Y/+^ tissues and in C57BL/6 FGFR2^+/+^ and C57BL/6 FGFRC342Y^+/+^ tissues. Gata1 is the 16^th^ top downregulated gene in C57BL/6 vs. BALB/c FGFR2^C342Y/+^ tissues (2.8-fold down in C57BL/6) and the top 15^th^ top downregulated gene in C57BL/6 FGFR2^+/+^ and C57BL/6 FGFR^C342Y+/+^ tissues (1.9-fold down in C57BL/6).

Analysis of the samples by the GO term osteoblast differentiation revealed 45/180 genes differentially expressed between C57BL/6 and BALB/c FGFR2^+/+^ tissues (ns), 40/180 genes between C57BL/6 and BALB/c FGFR2^C342Y/+^ tissues (p<.002), 35/182 genes between C57BL/6 FGFR2^+/+^ and C57BL/6 FGFRC342Y^+/+^ tissues (p<.001), and 3/178 genes between BALB/c FGFR2^+/+^ and BALB/c FGFRC^342Y+/+^ tissues (p not significant). Looking specifically at commonly used markers of osteoblast proliferation, the most consistent pattern to emerge is that expression of the FGFR^C342Y^ mutant receptor upregulates expression of osteoblast marker genes (*Spp1, Bglap, Runx2, Sp7* and *Alpl*) in C57BL/6 but not in BALB/c mice. *Runx2, Sp7* and *Alpl* are also expressed to a significantly greater extent in C57BL/6 vs. BALB/c FGFR2^C342Y/+^ mice.

### Confirmation of Candidate Genes by Real Time PCR

To confirm the RNA sequencing results, we next performed real time PCR on the top differentially regulated genes from our RNA sequencing analyses. Padi2, a histone H3 citrullination enzyme, is expressed at significantly higher levels in tissues from C57BL/6 FGFR2^+/+^ and FGFR2^C342Y/+^ mice than in their BALB/c counterparts. The proliferation promoting transcription factors Gata1 and E2F2 are expressed significantly less in in tissues from C57BL/6 FGFR2^+/+^ and FGFR2^C342Y/+^ mice than in their BALB/c counterparts. In addition, the de-ubiquitinase Uchl1 is expression significantly higher levels in tissues from C57BL/6 FGFR2^+/+^ and FGFR2^C342Y/+^ mice than in their BALB/c counterparts.

## Discussion

Activation mutations in *FGFR2IIIc* cause craniosynostosis of varying severity in humans and mice. As an initial step towards identifying potential genetic modifiers of the disorder, we backcrossed the FGFR2C^342Y/+^ mouse model of Crouzon syndrome onto inbred background strains. We found that FGFR2C^342Y/+^ mice on the C57BL/6 strain exhibits earlier onset and more extensive coronal suture fusion, worse skull shape abnormalities and a greater incidence of malocclusion than FGFR2C^342Y/+^ mice on the BALB/c strain. This data indicates increased craniofacial phenotype severity on the C57BL/6 background, which is consistent with other prior investigations which showed a more severe phenotype in the FGFR3^P244R^ mouse model of Meunke craniosynostosis when on the C57BL/6 as compared to the BALB/c strain (Twigg et al., 2009). We also recently noted a similar background dependent change in craniofacial phenotype severity in the *Alpl*^-/-^ mouse model of hypophosphatasia (data not shown). Together these findings indicate that comparison of C57BL/6 with BALB/c FGFR2C^342Y/+^ Crouzon mice may reveal genetic influences of craniosynostosis that are applicable to multiple forms of craniosynostosis.

We next performed RNA sequencing of cranial tissues isolated from C57BL/6 and BALB/c FGFR2C^342Y/+^ and FGFR2^+/+^ neonatal mice to identify potential modifying and mediator genes for severity of the Crouzon craniofacial phenotype and craniosynostosis. Initial analysis of the data demonstrated that substantially fewer genes were differentially expressed between BALB/c FGFR2^C342Y/+^ and FGFR2^+/+^ tissues than between C57BL/6 FGFR2^C342Y/+^ and FGFR2^+/+^ tissues, indicating that expression of the FGFR2C^342Y^ mutant receptor has a substantial influence on gene expression in cranial tissues of C57BL/6 mice but little influence on gene expression in cranial tissues of the BALB/c mice.

Analysis of the data by GO term craniosynostosis in addition to other genes known to be associated with coronal craniosynostosis, showed significantly diminished expression of *Msx1, Msx2*, and *Zic1* in C57BL/6 compared to BALB/c FGFR2^+/+^ tissues and diminished expression of *Msx1* and *Zic1* in C57BL/6 as compared to BALB/c FGFR2^C342Y/+^ mice. Because craniosynostosis-associated mutations in *Msx1, Msx2* and *Zic1* are gain of function (Liu et al., 1999; Twigg et al., 2015), and because we found downregulated expression of these genes in mutant mice on the more severe C57BL/6 background, these genes are unlikely to promote phenotype severity in FGFR2^C342Y/+^ mice, unless either too much or too little Msx1/Msx2 or Zic1 expression promotes craniosynostosis severity. The data also showed significantly diminished expression of *Twist1* in C57BL/6 compared to BALB/c FGFR2^+/+^ tissues. Craniosynostosis-associated mutations in *Twist* transcription factors are loss of function (el Ghouzzi et al., 1997; Yoshida et al., 2005) such that lower levels of *Twist* in C57BL/6 mice could predispose to a more severe craniosynostosis phenotype.

We also found significantly diminished expression of *En1* in C57BL/6 compared to BALB/c FGFR2^+/+^ and FGFR2^C342Y/+^ cranial tissues. *En1* is a homeodomain transcription factor that is essential for proper mesoderm/neural crest boundary formation of the coronal suture, and also for preventing premature osteoblast lineage commitment of suture progenitor cells (Deckelbaum et al., 2012). That *En*1 expression is down in C57BL/6 compared to BALB/c FGFR2^+/+^ and FGFR2^C342Y/+^ cranial tissues, suggests that low En1 expression levels may predispose to a more severe coronal suture fusion phenotype due to coronal suture boundary defects and/or a greater tendency for cranial progenitor cells to undergo osteoblast differentiation, consistent with its known function. It is also worth noting that *Twist* genes are in the same pathway as *En1*, eph/ephrin genes, *Msx2* and *FGFR2* in craniofacial bone and suture development (Deckelbaum et al., 2012; Merrill et al., 2006; Ting et al., 2009), such that this group of genes, when abnormally expressed, may represent an essential developmental pathway to premature coronal suture fusion. Differential expression of multiple members of the eph/ephrin gene family were found yet, no differential expression of *Efnb1* or *EphA4* members of the eph/ephrin gene family was found and these genes when mutated are established as causal for craniosynostosis in the form of coronal suture fusion (Ting et al., 2009; Twigg et al., 2004). It is important to note that this study utilized a neonatal developmental time point while previous studies investigating the expression and function of Efnb1 and Eph4a were performed at earlier embryonic time points. It is possible that differential expression for *Efnb1* and/or *EphA4* would be evident between C57BL/6 and BALB/c mice at an earlier developmental time point.

Of relevance to craniosynostosis, the epigenetic landscape controls cranial progenitor cell proliferation vs. osteoblast differentiation and osteogenesis (Dudakovic et al., 2015a; Dudakovic et al., 2018; Khani et al., 2017). The H3K27 methyl transferase Ezh2 was significantly downregulated in C57BL/6 compared to BALB/c FGFR2^342Y/+^ and in C57BL/6 FGFR2^+/+^ vs. FGFR2^342Y/+^ cranial tissues, indicating that diminished Ezh2 could be a primary mediator of increased phenotype severity in C57BL/6 FGFR2^342Y/+^ mice. Genetic ablation of Ezh2 leads to lack of craniofacial bone and cartilage formation in mice (Schwarz et al., 2014), while Prx1 conditional knockout of Ezh2 causes coronal craniosynostosis (Dudakovic et al., 2015b). Downregulation of Ezh2 gene expression in C57BL/6 compared to BALB/c FGFR2^342Y/+^ cranial tissues could drive cranial progenitor cells to preferentially differentiate over proliferate, which could lead to premature loss of cranial suture tissue and craniosynostosis. This hypothesis is also consistent with the fact that Wnt1-Cre driven Ezh2 conditional knockout mice exhibit similar shape abnormalities and craniosynostosis as that seen in FGFR2^342Y/+^ mice (Dudakovic et al., 2015b).

A principal component analysis of epigenetic regulators revealed that epigenetic regulator genes cluster differently dependent upon genotype and background strain. Analysis of the data by the GO term histone modifier genes revealed 13 genes that were differentially expressed between genotypes and background strains. Analysis by the GO term nucleosome genes revealed that six nucleosome genes were differentially expressed between C57BL/6 FGFR2^+/+^ and FGFR2^C342Y/+^ cranial tissues but none were differentially expressed between BALB/c FGFR2^+/+^ and FGFR2^C342Y/+^ cranial tissues. Multiple nucleosome genes were also differentially expressed between C57BL/6 and BALB/c of mutant and wild type genotypes. This data demonstrates that the epigenetic landscape is very likely different in cranial tissues of C57BL/6 and BALB/c mice, such that epigenetic landscape differences may predispose to a more severe craniosynostosis phenotype and may also be a primary mediator downstream of the FGFR2^342Y^ mutant receptor to exacerbate the phenotype in C57BL/6 mice. It should be noted that Padi2, a histone H3-R26 citrullination enzyme, is within the top 45 upregulated genes when comparing C57BL/6 with BALB/c FGFR2^+/+^ mice, making it a top phenotype modifier candidate gene.

The most consistent and striking finding of the RNA sequencing data was that minimal gene expression changes were induced upon expression of mutant FGFR2^342Y^ in BALB/c mice while numerous changes were induced upon expression in C57BL/6 mice. This pattern suggests the possibility that the FGFR2^342Y^ mutant receptor is either not expressed to the same extent in BALB/c as compared to C57BL/6 mice, and/or that the mutant receptor does not signal to induce cellular changes to the same extent in BALB/c as compared to C57BL/6 tissues. This data, combined with the fact that an analogous C278F Crouzon syndrome mutation leads to immature N-glycosylation of the receptor with increased trafficking for degradation in some cell types (Hatch et al., 2006), led us to hypothesize that craniosynostosis severity in Crouzon syndrome may be mediated by preferential mutant receptor trafficking to membranes for signaling vs. to the lysosome for degradation that is dependent upon genetic background. Results from this RNA sequencing analysis support this hypothesis. The data showed increased expression of N-glycosylation enzymes and ER/Golgi chaperone proteins *Hspa1a* and *Hspa1b* in C57BL/6 vs. BALB/c FGFR2^+/+^ tissues and in C57BL/6 vs. BALB/c FGFR2^342Y/+^ cranial tissues, which supports the concept that the difficult to process mutant FGFR2^342Y^ protein may have a greater chance of being retained in the ER/Golgi for N-glycosylation and processing in the C57BL/6 than the BALB/c mice. Though it must also be noted that *Hspa1a* and *Hspa1b* genes were also upregulated in BALB/c FGFR2^+/+^ vs. FGFR2^342Y/+^ cranial tissues, while chaperones Clgn and Hsbp were downregulated in C57BL/6 FGFR2^+/+^ vs. FGFR2^342Y/+^ tissues, suggesting that the mutant receptor likely feeds back on chaperone proteins differently in the two strains of mice. Analysis for the GO term lysosome revealed that expression of FGFR2^342Y^ increased expression of glycoside trimming enzymes and decreased expression of peroxides and proteases only in the C57BL/6 mice. This data is perhaps indicative that the mutant receptor is more likely to complete maturation of glycosylation (involves trimming) and of a lesser tendency for lysosomal protein degradation in the C57BL/6 mice. Finally, analysis of top differentially expressed genes revealed that the *Uchl1* deubiquitinating enzyme, was upregulated by 3-fold in C57BL/6 over BALBc FGFR2^+/+^ and 2.6-fold in C57BL/6 over BALBc FGFR2^C342Y/+^ cranial tissues. *Uchl1* was the 21^st^ top upregulated gene between C57BL/6 and BALB/c FGFR2^+/+^ mice, making it a strong potential phenotype modifier candidate gene.

Real time PCR results were largely consistent with the RNA sequencing results. *Padi2* and *Uchl1* were expressed to a significantly greater extent in C57LB/6 than BALB/c FGFR2^+/+^ and FGFR2^C342Y/+^ tissues, while *Gata1* and *E2F2* were expressed to a significantly greater extent BALB/c FGFR2^+/+^ and FGFR2^C342Y/+^ than C57LB/6 tissues when assayed by real time PCR and RNA sequencing. That said, the real time PCR results did not show a significant difference between C57BL/6 FGFR2^+/+^ and FGFR2^C342Y/+^ mRNA levels while that was found in the RNA sequencing results. This could be due to the variation seen in the C57BL/6 FGFR2^+/+^ mRNA levels as assessed by real time PCR. In addition, higher levels of Gata1 mRNA were seen in BALB/c FGFR2^C342Y/+^ than BALB/c FGFR2^+/+^ tissues, while only a trend was seen in the RNA sequencing data.

The idea that “activating” mutations in genes can lead to increased degradation of the mutant protein in certain cell types or environments is not new. Somatic mutations in EGFRs are known to influence prognosis of lung adenocarcinoma and also increase degradation of the mutant receptor (Chung et al., 2016). The Boston-type craniosynostosis “activating” mutation in Msx2 was previously shown to increase degradation of the receptor (Yoon et al., 2008). Additionally, *FGFR2IIIc* expression is not limited to cranial tissues but the Crouzon syndrome phenotype is primarily craniofacial. *FGFR2IIIc* is highly expressed in other mesenchymal tissues including smooth muscle tissue of the stomach and gastrointestinal tract (Hughes, 1997). In addition, overexpression of *FGFR2IIIc* and/or somatic mutations that are identical to those seen in the FGFR2-associated craniosynostosis syndromes are associated with gastric cancer and poor prognosis (Hattori et al., 1996; Pollock et al., 2007), yet individuals carrying craniosynostosis syndrome associated mutations in FGFR2 do not to our knowledge have an increased risk of gastric or other cancers. Together, these findings indicate that craniosynostosis-associated mutations in FGFR2 lead to diminished mutant receptor protein expression in some cell types. The RNA sequencing data presented here indicates that the FGFR2^C342Y^ mutation likely leads to altered protein processing and diminished mutant protein expression in the BALB/c as compared to C57BL/6 genetic background. In future studies we will work to determine if strategies developed to degrade target proteins in cancer can be used to decrease protein expression of FGFR2^C342Y^ to diminish craniosynostosis phenotype severity (Chung et al., 2016; Naito et al., 2019).

## Conclusions

Together, the data presented here demonstrates that the Crouzon craniosynostosis phenotype is genetic background dependent and that C56BL/6 mice exhibit a more severe phenotype than BALB/c mice upon expression of Crouzon craniosynostosis-associated mutant FGFR2^C342Y^. C56BL/6 mice exhibit greater differential gene expression than BALB/c mice upon expression of mutant FGFR2^C342Y^. More specifically, C56BL/6 mice exhibit differential expression of genes associated with tissue boundaries, cellular proliferation, osteoblast differentiation, epigenetics, ER/Golgi protein processing and lysosome components upon expression of mutant FGFR2^C342Y^. Differential expression of protein processing and degradation genes indicates that the milder BALB/c phenotype may be due to increased mutant protein degradation in BALB/c mice, a hypothesis we will pursue further in future studies.

## Author Contributions

**Hwa Kyung Nam:** Methodology, Investigation, Investigation, Validation, Writing – Review&Editing. **Amel Dudacovik.**: Methodology, Investigation, Data curation, Software, Investigation, Validation, Funding Acquisition, Writing – Review&Editing. **Andre VanWijnen**: Conceptualization, Resources, Methodology, Investigation, Visualization, Supervision, Project Administration, Funding Acquisition, Writing – Reviewing and Editing. **Nan Hatch**: Conceptualization, Methodology, Resources, Investigation, Visualization, Supervision, Project Administration, Funding Acquisition, Writing-Original draft preparation.

## Acknowledgements

The authors would like to thank Dana M. King for her bioinformatic support of RNA sequencing meta-analyses (Bioinformatics Core, University of Michigan). This work was supported by grant R01DE025827 from NIDCR (to N.E.H.) grant P30AR069620 from NIHMS, grant R01AR049069 from NIAMS (to AJvW) and a Career Development Award in Orthopedics Research (to A.D.) from the Mayo Clinic, Rochester MN.

## Abbreviations

FGFR: Fibroblast Growth Factor Receptor
Msx1: Msh homeobox 1
Msx2: Msh homeobox 2
Twist1: Twist-related protein 1
Zic1: Zinc finger of the *c*erebellum 1
En1: Homeobox protein engrailed-1
Ezh2: Enhancer of zeste homolog 2
Mgat5b: Alpha-1,6-mannosylglycoprotein 6-beta-N-acetylglucosaminyltransferase B
Mgat5: Alpha-1,6-mannosylglycoprotein 6-beta-N-acetylglucosaminyltransferase A
Hspalb: Heat shock 7OkDa protein 1B
Hspa1a: Heat shock 70 kDa protein 1
Hspa2: Heat shock-related 70 kDa protein 2
Clgn: Calmegin
Hspb1: Heat shock protein 27
Hexa: Hexosaminidase A
Hexb: Hexosaminidase B
Mpo: Myeloperoxidase
Prtn2: Proline-rich transmembrane protein 2
Prss57: Serine Protease 57
Uchl1: Ubiquitin carboxy-terminal hydrolase L1
Mik67: Proliferation-Related Ki-67 Antigen
Ccnb1: Cyclin B1
Aurkb: Aurora kinase B
Cdk1: Cyclin-dependent kinase 1
Ccnb2: Cyclin B2
Gata1: GATA-binding factor 1
E2F2: E2F Transcription Factor 2
Spp1: Osteopontin
Bglap: Bone gamma-carboxyglutamic acidcontaining protein
Runx2: Runt-related transcription factor 2
Sp7: Osterix
Alpl: Tissue-nonspecific alkaline phosphatase
Padi2: Protein-arginine deiminase type-2
Efn: Ephrin
Eph: Eph receptor

---

**Supplemental Figure 1.**
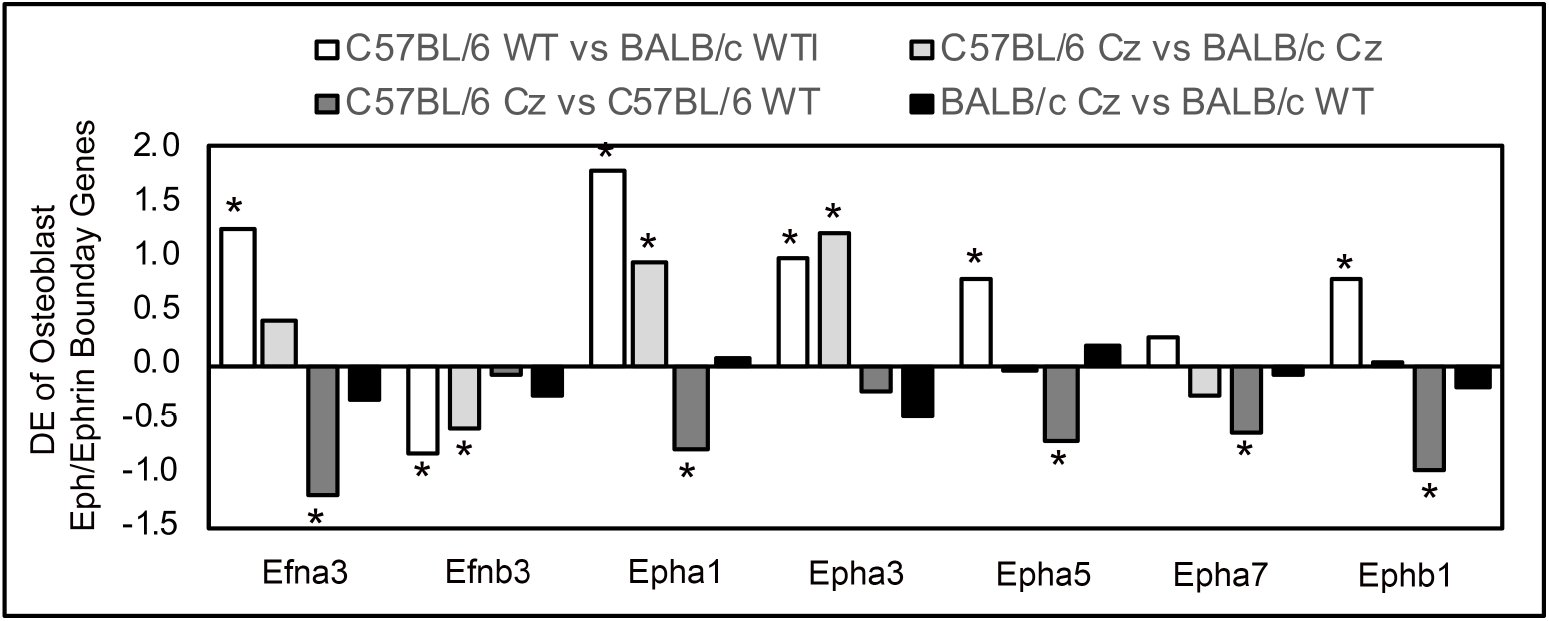
Differential Expression of Eph/Ephrin (Developmental Boundary) Genes. All differentially expressed eph/ephrin genes are shown. Data for each gene is presented for four different individual comparisons, as indicated by color.

**Supplemental Figure S2.**
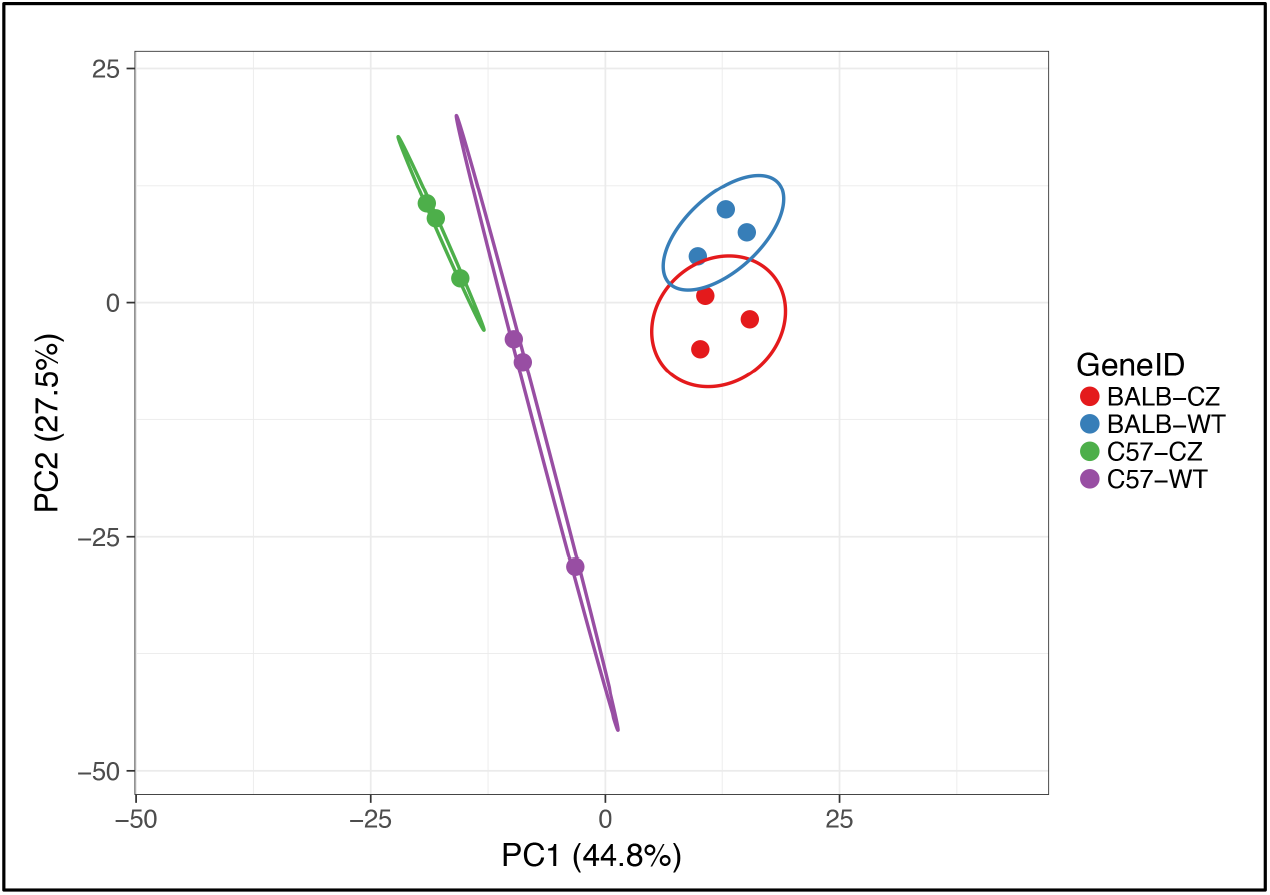
Principal Component Analysis of Epigenetic Regulators. A principal component analysis of epigenetic regulators including principal component 1 (27.5% of differences) and 2 (44.8% of differences) demonstrates segregation of epigenetic regulator gene expression by genotype and genetic strain.

**Supplemental Table 1.** Differential Expression of Histone Modifying Genes. Log fold change (Log Fc) and Bonferroni adjusted p values for differential expression of the GO term histone modifier genes is shown for each of the individual comparisons.

| Gene Name | Gene Function | Log Fc<br>C57BL/6 WT<br>vs.<br>BALB/c WT | Adj<br>p value | Log Fc<br>C57BL/6 Cz<br>vs.<br>BALB/c Cz | Adj<br>p value | Log Fc<br>C57BL/6 Cz<br>vs.<br>C57BL/6 WT | Adj<br>p value | Log Fc<br>BALB/c Cz<br>vs.<br>BALB/c WT | Adj<br>p value |
| --- | --- | --- | --- | --- | --- | --- | --- | --- | --- |
| Eya1 | histone<br>dephosphorylation | 0.83 | 1.00E-06 |  |  | -0.82 | 1.00E-06 |  |  |
| Eya2 | histone<br>dephosphorylation |  |  |  |  | -0.94 | 1.00E-06 |  |  |
| Eya4 | histone<br>dephosphorylation | 1.72 | 1.00E-06 | 0.91 | 9.96E-05 | -1.03 | 1.00E-06 |  |  |
| Padi2 | histone H3-R26<br>citrullination | 2.96 | 1.00E-06 | 1.36 | 1.00E-06 | -1.06 | 1.00E-06 |  |  |
| Padi4 | histone H3-R26<br>citrullination |  |  | -0.91 | 7.63E-03 |  |  |  |  |
| Drd1a | histone-serine<br>phosphorylation |  |  |  |  | 0.65 | 8.84E-03 |  |  |
| Ccnb1 | histone-serine<br>phosphorylation |  |  | -0.75 | 1.00E-06 | -0.68 | 1.00E-06 |  |  |
| Hdac4 | histone<br>deacetylase | 0.93 | 1.00E-06 | 0.93 | 1.00E-06 | 0.24 | 1.14E-01 |  |  |
| Hdac9 | histone<br>deacetylase | 0.63 | 7.11E-04 | 0.99 | 1.00E-06 | 0.31 | 5.80E-02 |  |  |
| Hdac11 | histone<br>deacetylase | 0.68 | 4.42E-04 | -0.10 | 7.12E-01 | -0.70 | 6.15E-05 |  |  |
| Prkaa2 | histone serine<br>kinase activity | 1.15 | 1.43E-04 |  |  | -0.98 | 3.38E-04 |  |  |
| Aurka | histone serine<br>kinase activity |  |  | -0.85 | 1.00E-06 | -0.61 | 1.00E-06 |  |  |
| Aurkb | histone-serine<br>phosphorylation,<br>histone serine<br>kinase activity |  |  | -0.86 | 1.00E-06 | -0.60 | 1.00E-06 |  |  |

**Supplementary Table 2.** Differential Expression of Nucleosome Genes. Log fold change (Log Fc) and Bonferroni adjusted p values for differential expression of the GO term nucleosome genes is shown for each of the individual comparisons.

| Gene Name | Log Fc<br>C57BL/6 WT<br>vs.<br>BalbC.WT<br>DE | Adj<br>p value | Log Fc<br>C57BL/6 Cz<br>vs.<br>BalbC.Cz<br>DE | Adj<br>p value | Log Fc<br>C57BL/6 Cz<br>vs.<br>C57BL/6 WT<br>DE | Adj<br>p value | Logfc<br>BalbC.Cz<br>vs.<br>BalbC.WT<br>DE | Adj<br>p value |
| --- | --- | --- | --- | --- | --- | --- | --- | --- |
| Hist2h2be | 2.23 | 0.000 | 0.98 | 0.0003 | -1.23 | 0.000 |  |  |
| Hist2h4 | 0.97 | 0.003 |  |  | -1.08 | 0.000 |  |  |
| Myst3 | 0.90 | 0.000 |  |  |  |  |  |  |
| Hist3h2a | 0.87 | 0.000 | 0.78 | 0.0000 |  |  |  |  |
| Hist1h4i | 0.73 | 0.005 |  |  | -0.62 | 0.006 |  |  |
| H1foo | 0.67 | 0.021 |  |  |  |  |  |  |
| Hist2h3c2 | 0.63 | 0.044 | 1.08 | 0.0003 |  |  |  |  |
| Hist1h4a | -0.64 | 0.038 |  |  |  |  |  |  |
| Hist1h3e | -0.65 | 0.022 |  |  |  |  |  |  |
| H2afx | -0.70 | 0.000 | -1.10 | 0.0000 |  |  |  |  |
| H2afj | -0.82 | 0.000 | -0.85 | 0.0000 |  |  |  |  |
| Hist1h3c | -0.91 | 0.006 | -1.12 | 0.0008 |  |  |  |  |
| Irf4 | -0.96 | 0.001 | -1.67 | 0.0000 |  |  |  |  |
| Hist2h2ac | -1.77 | 0.000 | -2.03 | 0.0000 |  |  |  |  |
| Cenpa |  |  | -0.88 | 0.0000 |  |  |  |  |
| Hist1h1e |  |  | -0.79 | 0.0225 | -0.71 | 0.016 |  |  |
| Myst4 |  |  |  |  |  |  |  |  |
| Hist1h1b |  |  | -0.87 | 0.0124 |  |  |  |  |
| Hist1h1a |  |  | -0.69 | 0.0482 |  |  |  |  |
| Hist1h3g |  |  | -1.12 | 0.0002 |  |  |  |  |
| Hist1h3f |  |  | -0.74 | 0.0185 |  |  |  |  |
| Hist1h3i |  |  | -0.76 | 0.0169 |  |  |  |  |
| Hist1h4d |  |  | -0.70 | 0.0443 |  |  |  |  |
| Hist1h4f |  |  | -0.92 | 0.0022 |  |  |  |  |
| Hist1h4j |  |  |  |  | -0.67 | 0.006 |  |  |
| Hist1h2ak |  |  | -0.70 | 0.0143 |  |  |  |  |
| Hist1h2bb |  |  |  |  | -0.63 | 0.006 |  |  |
| Hist1h3a |  |  | -0.92 | 0.0056 |  |  |  |  |
| Hist1h4m |  |  | -1.07 | 0.0008 |  |  |  |  |

